# Guidance of circumnutation of climbing bean stems: An ecological exploration

**DOI:** 10.1101/122358

**Authors:** Paco Calvo, Vicente Raja, David N. Lee

## Abstract

In this report we explore the guidance of circumnutation of climbing bean stems under the light of general rho/tau theory, a theory that aims to explain how living organisms guide goal-directed movements ecologically. We present some preliminary results on the control of circumnutation by climbing beans, and explore the possibility that the power of movement in plants, more generally, is controlled under ecological principles.

## Introduction

Living organisms guide the movement of effector organs or cells in response to stimuli to make, or avoid, contact with things—be it a bee flitting from flower to flower, a gibbon swinging from branch to branch, or a peregrine falcon diving on a flying pigeon. Strikingly enough, plants, unlike animals, are commonly believed to remain still, with their behavioural repertoire reducing, or so the story goes, to invariant tropistic—Jacques Loeb’s (1918) ‘forced reactions’—or nastic responses implemented in the form of sets of fixed reflexes. The need to control their movements is thereby eliminated or seriously undermined. And yet plants are as much in the move as any other living organism. Plant stems grow alternatively on different sides, which results in the stem bending in one direction, then in the opposite one. But there is virtually no growing part of any single plant that fails to exhibit a movement of nutation (Mugnai et al., 2007). Not only the tips of shoots sway in circles as they grow, but also leaves and roots exhibit ‘revolving nutation’, as Julius von Sachs called it, or *circumnutation*, to use the expression coined by Charles Darwin. Circumnutation, we may say, is universal. All plants do it. Shoots of climbing plants guide their movements to reach a support; roots navigate belowground, guiding their movements to secure nutrients intake; young and terminal leaves display helical and rotational oscillatory movements, etc. (Darwin, 1975; Darwin and Darwin, 1880).

In this report, we consider the possibility that *the power of movement in plants*, to echo Darwin and his son’s seminal work, is not forced, or hardwired, but rather appropriately controlled as much as the movements performed by bees, gibbons or peregrine falcons are. More specifically, we shall focus our attention on what is probably the simplest form in which general circumnutation can be modified: the one exhibited by twining plants, in particular, by common beans (*Phaseolus vulgaris*) as they approach and twine spirally round supports for photosynthetic purposes. Unlike leaf-climbers, tendril-bearers, and hook and root climbers that use a whole new bag of tricks insofar as attachment mechanisms and stem structure and function are concerned (Isnard & Silk, 2009), bean shoots rely exclusively on an increase in the amplitude of an otherwise ordinary movement of circumnutation. Such basic, and yet modified, revolving nutation shall be the focus of our attention. In particular, we aim to explore the guidance of circumnutation of climbing bean stems under, broadly speaking, Gibsonian ecological principles (Gibson, 1966; 1979); and more specifically, under the light of General Tau Theory (Lee 1998; Lee et al., 2009).

## The control of movement in climbing plants

The underlying idea that motivates the research herewith reported is the suspicion that the control of movement in plants is not unlike the control of movement in animals. Plants and animals, we contend, have functionally similar internal systems for organizing sets of behaviours. In essence, a plant that orients towards, say, a source of energy behaves in functionally the same way as an animal that runs towards its prey. It is in this sense that the type of control required to perform such actions is our object of study.

Nutation is due to differential cell growth, and not to changes in the state of turgidity (rigidity) of cells, as is the case, for instance, in heliotropic and nyctinastic (sleep) movements. Whereas the latter exploit changes in turgor pressure and are thus reversible by the alternative gain and loss of cell water, the former, being dependent upon growth, is irreversible. In addition, growth-related circumnutation of the stem is not triggered by external forces themselves, such as temperature, gravity, or day/night cycles, but is rather brought about, maintained and modified by endogenous means. Plants explore, and exploration uses up energy and therefore needs to be done efficiently, especially considering that growth-related movements are irreversible. Control thus appears to be needed for the regular pattern of bending observed to obtain. In particular, both the direction and the amplitude of nutational movements require control, if the metabolic cost of irreversible but idle movements is to be minimized.

With that being said, that plants or animals *control* their movements does not imply that their behaviour is to be accounted for in computational or information-processing terms. In fact, our working hypothesis is that both plants and animals guide their movements *ecologically*—non-computationally. According to ecological psychology, plants, like animals, perceive what is available in terms of biologically relevant interactions (Carello et al., 2014). Plants perceive opportunities for behavioural interaction with their local environment in the form of what Gibson dubbed “affordances”. Climbing plants are in this way ecological *perceivers*. Vines perceive possibilities for action, such as when a support is perceived as affording climbing. Or take *Monstera gigantea*, a climbing vine whose seeds are able to perceive an affordance (climb-ability) skototropically, as they grow towards darkness (Turvey et al., 1981).

Under this framework, the proper unit of analysis is the whole organism-environment system as such (Richardson et al., 2008). A climbing plant and its support constitute an ecologically coupled system in which the action of twining and the perception of affordances form a continuous and cyclic loop. Despite things being in constant flux, some relations remain unchanged, and organisms can pick them up. This information is relational, and takes the form of invariant properties of the underlying structure of an ever-changing environment that can in principle be directly detected. Ecological psychologists say that environmental information is *specificational:* information in the vicinity of a climber specifies ways for the plant to interact with features, such as the support standing nearby. Our working hypothesis is that plants, like animals, pick up the invariant structure of an ever-changing environment. General rho/tau theory puts some flesh onto this framework for empirical test.

## General Rho/Tau Theory

*General rho/tau theory* (Lee 1998; Lee et al. 2009) aims to explain how living organisms guide goal-directed movements endogenously by using prescriptive and perceptual information. Up to 2009, the theory dealt with guidance of movement in animals (see Lee et al., 2009). In this section we review its main tenets, and elaborate on how the theory applies to plants too. In a nutshell, the main points of general rho/tau theory are as follows:

i. Purposeful, goal-directed, movement entails guiding the trajectory of an effector to a goal across a motion-gap; that is, it requires the guidance of the closure of motion-gaps, where a motion-gap is defined as the changing gap between a current state and a goal state. Motion-gaps may occur across a variety of dimensions—e.g., distance when reaching, angle when shifting gaze or direction of movement, pressure when gripping, pitch and loudness when vocalizing or making a noise, intraoral pressure when suckling, etc.
ii. Closing a motion-gap requires:

a. Generating prescribing information to specify the intended trajectory of the gap;
b. picking up perceptual information from the stimulus about how the trajectory is actually evolving; and
c. regulating the motor information to make the prescribing and perceptual information match.
iii. The primary perceptual information used in guiding a goal-directed movement is transmitted through the medium of *power* (rate of flow of energy), and takes the form of the *rho/tau of a power gap* (the tau of a gap of magnitude, *X*, is the time it will take the gap to close at the current closure rate). The tau of the gap equals the current magnitude, *X*, of the gap divided by the current rate of change of *X*, viz. 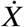. The defining equation is:

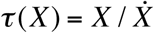
The unit of *tau* is time. Since rho and tau are mathematical duals—rho of a gap = 1/tau of the gap—, for ease of exposition, rho rather than tau of a gap is used here. We shall then say that the information used for guiding the closing (or opening) of a motion-gap is the rho of the gap, the proportionate rate of closing (or opening) of the gap. Rho of gap X at any time t equals the current rate of change of the size of the gap divided by the current size of the gap. Thus,

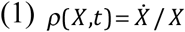
where the dot indicates the time derivative. Rho of a motion-gap is, in principle, directly perceptible by all known perceptual systems: in contrast, the size of a motion-gap, or any of its time derivatives (velocity, acceleration etc), is not directly perceptible or specified in the stimulus (Lee, 1998), requiring a scaling factor.
iv. Synchronizing the closing of two gaps, as when catching a ball, is achieved by rho-coupling the gaps, by keeping the rho of one gap (e.g., hand to catching place) in constant proportion with the rho of another gap (e.g., ball to catching place). Hence, the general rho-coupling equation

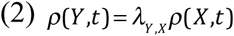
where Y and X are the gaps, t is time and *λ_y,x_* is the coupling factor, which determines the shape of the velocity profile of Y relative to X.
v. By rho-coupling the motion-gap (via perceptual information) onto a changing ‘guiding-gap’ generated endogenously, prescribes how a motion-gap, Y, should close. Analyses of many skilled movements indicate a guiding-gap, G, that changes at a constant accelerating rate from rest. Thus, an extrinsic motion-gap, Y, is guided by making the movement follow the equation

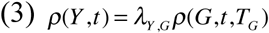
where *T_G_* is the time the guiding gap G takes to close (or open). Here

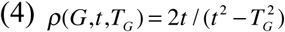
as derived from Newton’s equations of motion, where time t runs from 0 to *T_G_*.^1^

Figures 1A-1D below depict the effect of the coupling factor, *λ_X,G_* on rhoG-guidance of gap X following the equation *ρ*(*X,t*) = *λ_X,G_* (*X,t,T,_G_*)^2^.

**Figure 1A.**
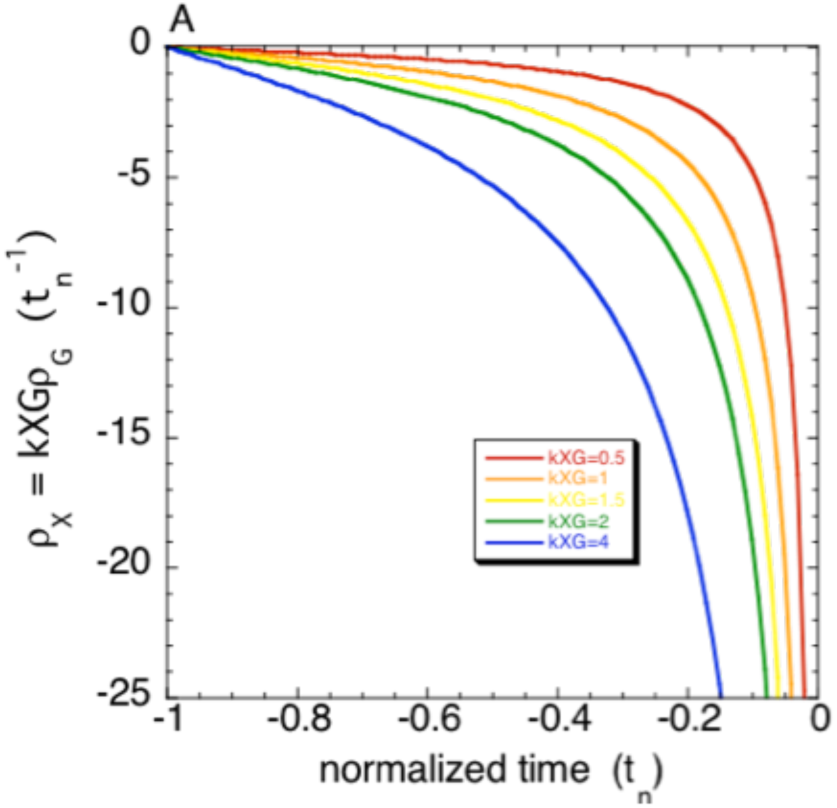
Effect of the coupling factor, *λ_X,G_* on rhoG-guidance of gap X following the equation *ρ*(*X,t*) **=** *λ_XG_*(X, *t,T*_G_). (A) *ρ_X_*, the rho of gap.

**Figure 1B.**
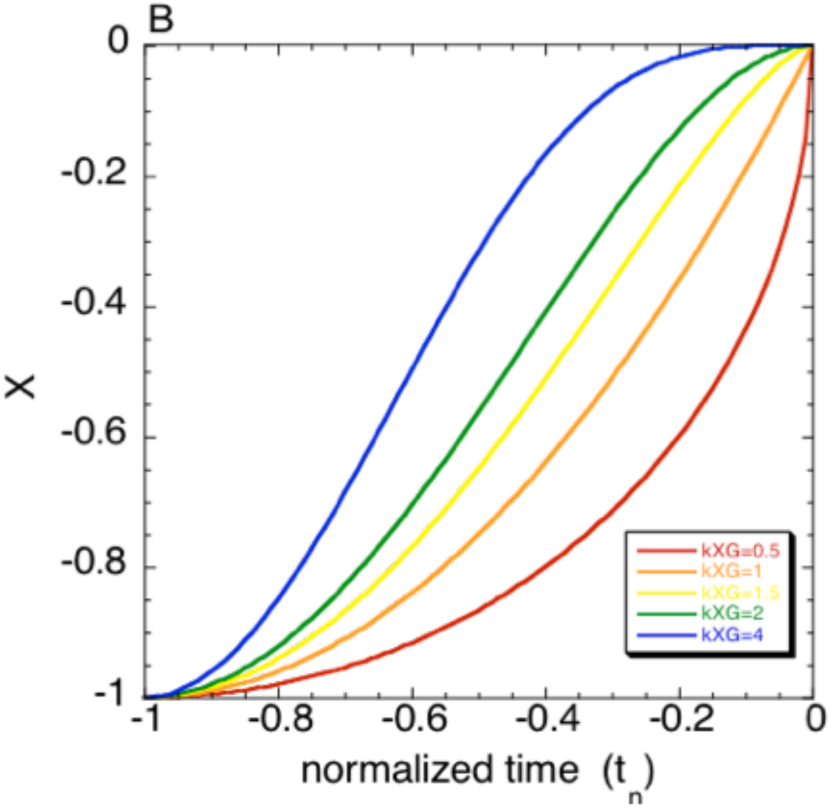
Effect of the coupling factor, λ_X,G_, on rhoG-guidance of gap X following the equation ρ(*X,t*)=λ(*X,t,T,_G_*). (B) X, the size of the gap.

A motion-gap that follows Eqs. (3) and (4) is said to be rhoG-guided. The kinematic form of the prescribed motion-gap is defined by Eqs. (3) & (4). There are two adjustable parameters,

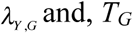
which specify, respectively, the shape of the velocity profile and the duration of the motion-gap. The velocity profile is single-peaked and the position of the peak is determined by the value of *λ_Y,G_* (Fig. 1C).

**Figure 1C.**
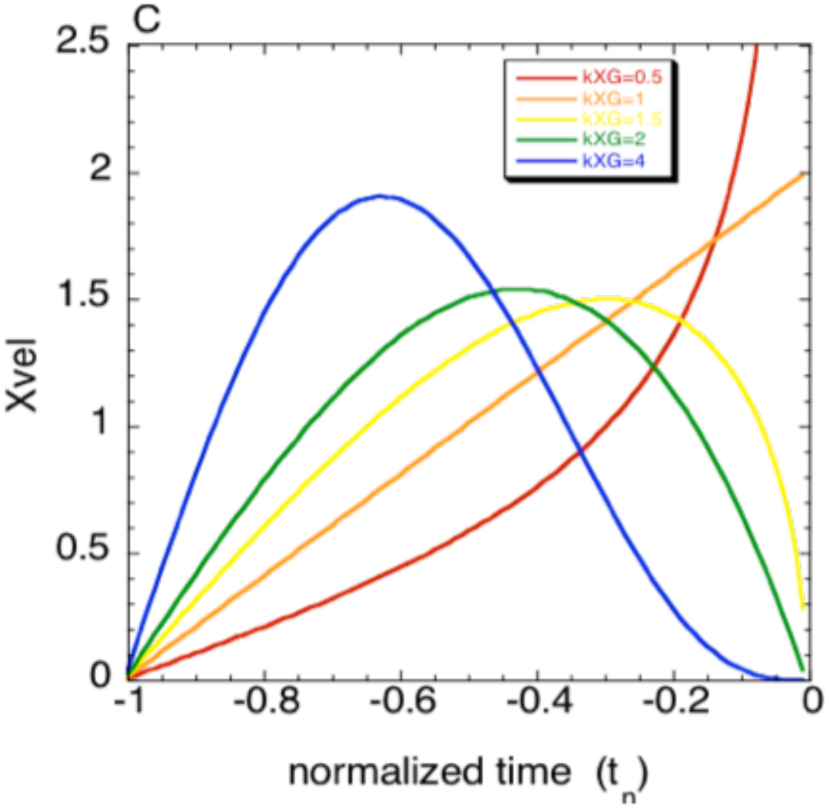
Effect of the coupling factor, λ_X,G_, on rhoG-guidance of gap X following the equation *ρ*(*X,t*) = *λ_χ G_*(*X,t,T_G_*). (C) Xvel, the velocity of closure of the gap.

When *λ_Y,G_* > 1 the gap-closing movement first accelerates at a varying rate up to a peak velocity and then immediately decelerates at a varying rate to the goal (Fig. 1D).

**Figure 1D.**
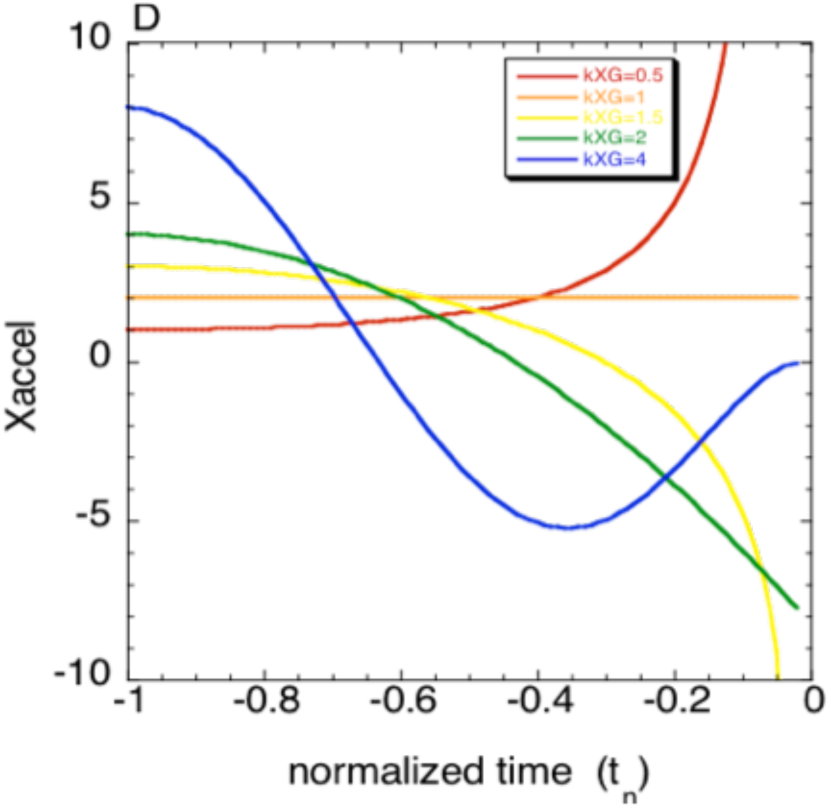
Effect of the coupling factor, λ*_X,G_* on rhoG-guidance of gap X following the equation *ρ*(*X,t*) = *λ_X,G_*(*X,t,T_G_*). (D) Xaccel, the acceleration of closure of the gap.

Gently touching an object, so that the velocity of approach is zero when the object is reached, requires *λ_Y,G_* ≥2. Hitting something, so that when the object is reached the velocity of approach is positive, requires *λ_Y,G_* <2. Thus, rhoG-guidance of motion-gaps is a simple way of regulating goal-directed movement.

General rho/tau theory has been tested successfully across a number of species and activities (Lee et al. 2009). High-resolution movement analysis has yielded evidence for rho/tauG-guidance of motion-gaps that span a range of skills, including newborn babies suckling (Craig & Lee 1999), infants catching (van der Meer et al. 1994), adults reaching (Lee et al. 1999), controlling gaze (Grealy et al. 1999; Lee 2005), intercepting (Lee et al. 2001), putting at golf (Craig et al., 2000), flying aircraft (Padfield 2011), singing and playing music (Schogler et al. 2008), and flies landing (Wagner 1982), hummingbirds feeding (Delafield-Butt et al. 2010). Also, evidence of rho has been found in the electrical activity in the brains of locusts (Rind & Simmons 1999), pigeons (Sun and Frost 1998), monkeys (Merchant et al. 2004) and humans (van der Weel et al. 2009), and in a unicellular paramecium (Delafield-Butt et. al 2012).

There are two basic types of movement: *propriospecific* movements that are specific to the individual’s body and *expropriospecific* movements that are specific to the organism’s relation to the environment and other organisms. In animals, there are vital propriospecific movements within the skeletal, muscular, respiratory, cardiovascular, lymph-vascular, endocrine, digestive, excretory, and reproductive systems. Similar life-sustaining propriospecific movements occur in plants, cells, and fungi. Vital expropriospecific movements in animals include moving in the environment, grasping objects, feeding, avoiding predators, and mating. Again, similar expropriospecific movements occur in plants, cells and fungi.

In all cases, propriospecific and expropriospecific movements must be coordinated to achieve functional movement. This is the essential task of the electrochemical nervous and endocrine informational systems in animals. These pick up, generate and integrate information for guiding movements. *Sensory* information about movement is picked up by active perceptual systems, both within the body and at its surface. *Prescriptive* information specifying purposive movement is generated within the electrochemical informational systems. Both types of information flow along channels in the body as *electrochemical power* (rate of flow of energy). The information is a mathematical rho/tau function of the electrochemical power. The prescriptive and sensory information are integrated in the nervous and endocrine systems, resulting in rho/tau *motor* information being transmitted to contractile cells in muscles in animals and, we believe, in roots and stems in plants. In animals, the basic function of the nervous and endocrine systems is to organize the prescriptive, sensory and motor information to achieve purposeful movement. Our working hypothesis is that the same applies in plants.

Like animals, plants need ways of transmitting the prescriptive, perceptual and muscular-like information around their bodies. Plants thus fall under the scope of general rho/tau Theory. A bean shoot, as it grows and seeks a support with its tip, circumnutates the whole shoot in its endeavour. Plants also need *perceptual* organs to seek out information for guiding their effectors. They also need to generate *prescribing* information to specify the movement trajectory necessary to fulfil their purpose.

## Behavioural study

The basic experimental paradigm in our study consisted in the analysis of plant circumnutation behaviour through controlled time-lapse observation of the approaching manoeuvre of a bean plant engaged in closing the gap in between its shoot apex and the support that stands in its vicinity.

## Method

### Subjects

The experiment was conducted on bean plants (*Phaseolus vulgaris, Leguminosae*). Healthy-looking bean seeds were selected from *Semillas Ramiro Arnedo* (Av. Infante D. Juan Manuel, ed. Cinco Estrellas, 30011, Murcia, Spain). Seeds were potted and kept at ambient temperature. Water levels were checked periodically to maintain a hydric bedding throughout the experiment.

### Apparatus

A potted bean plant was placed at the centre of the experimental growing chamber. A single vertical pole (not shown in figures 1 and 2), placed at a distance of 50 centimetres from the plant centre was used as potential support for the bean to twine around. Two digital single-lens webcams, one (fig. 1) at a height of 200 cms. pointing vertically down with the X-axis of the picture frame closely parallel to the line joining the centre of the plant (C) and the bottom of the pole; the other camera (fig. 2) located at 50 cms. from the plant, pointing horizontally with the X-axis of its picture frame approximately horizontal.

**Figure 1.**
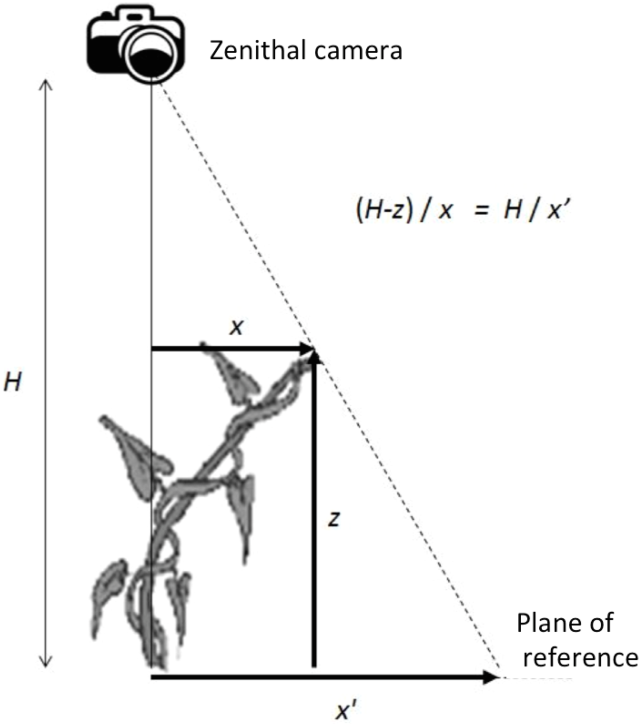
Zenithal camera. An illustration of the experimental apparatus.

**Figure 2.**
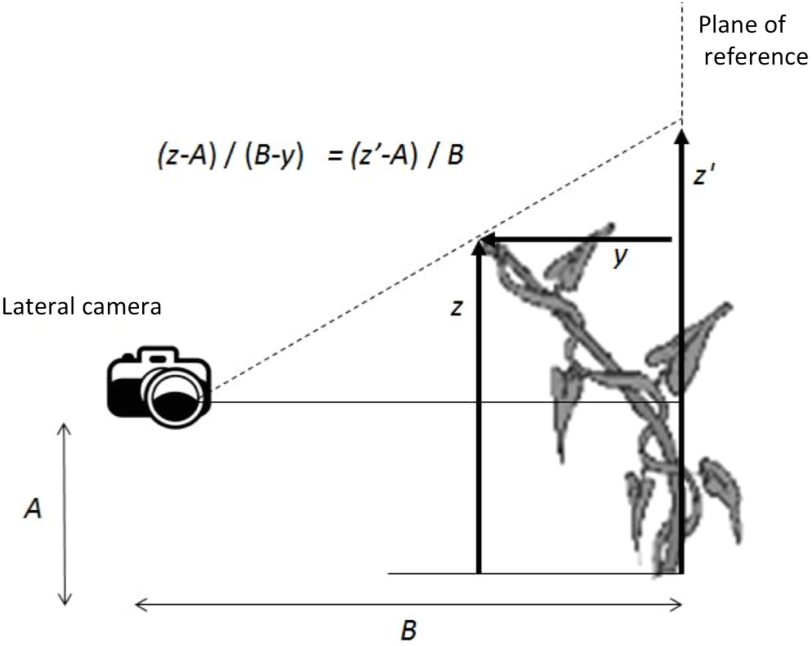
Lateral camera. An illustration of the experimental apparatus.

## Procedure

The setting was recorded from the two view-points illustrated in figures 1 and 2. Lenses were zoomed to optimize the resolution of the movement of the shoot apex.

From the camera recordings, the (x,y,z) coordinates of the plant’s growing tip, T; the bottom of the pole, P; and the centre of the plant, C, were computed—where the z-axis was vertical, the (x,y) plane horizontal and the x-axis approximately parallel with the line between P and C. Time-lapse records were made, the time interval between frames being 30 seconds. The pole was vertical and in place during the whole recording session.

We analysed the movement of the shoot-tip by digitizing the time-lapse frames. To do this, we first recorded the coordinates of the shoot base, the shoot tip, and the pole by digitizing those three points on one single picture frame. Subsequently, we traced the movement of the shoot apex by digitizing the coordinates of the shoot tip in each of the 1,034 time-lapse frames that had been recorded (both zenithal and lateral), as the plant moved throughout the experiment. This series of digitized points gave us a file with the 2-D coordinates of the shoot apex for each zenithal and lateral picture being taken. Out of these zenithal and lateral coordinates, we obtained a file with the (x’, y’, z’) coordinates of the whole 3-D setting.

Because optical aberrations had to be dealt with before any further analysis could be made on coordinates (x’, y’, z’), perspective error was measured.^3^ Distorted coordinates (x’, y’, z’) were transformed into coordinates (x, y, z), by:

i. Digitizing distorted coordinates (x’, y’, z’) for both cameras (see above).
ii. Measuring distances H, A and B (figures 1 and 2, above, respectively)—A = 7 cm y B = 50 cm, y H=125 cm.
iii. Calculating real, undistorted coordinate z out of y’ following equation:

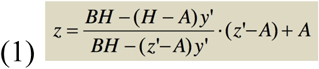
iv. Calculating real, undistorted coordinates x and y, out of z, following equations:

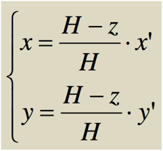

The (x,y) coordinates for the shoot tip were subsequently transformed into (r,A) coordinates (the zenithal camera allowed us to track the (r, A) coordinates of the shoot tip, where *r* is the horizontal radial distance of the tip from the pole, and *A* is the angle of this radius vector from a horizontal reference direction fixed in this environment). We then performed a rhoG/tauG analysis on these (r,A) coordinates.

## Results

The following analysis is of circumnutation in the presence of the pole. In figure A, shoot tip, T, appears to circumnutate around an approximately elliptical orbit, centred on the mean (x,y) position, G, of the tip. The major axis of the orbit points roughly in the general direction of the pole, P. There does not appear any systematic change over time with the orbit, except for the blue orbit.

**Figure A.**
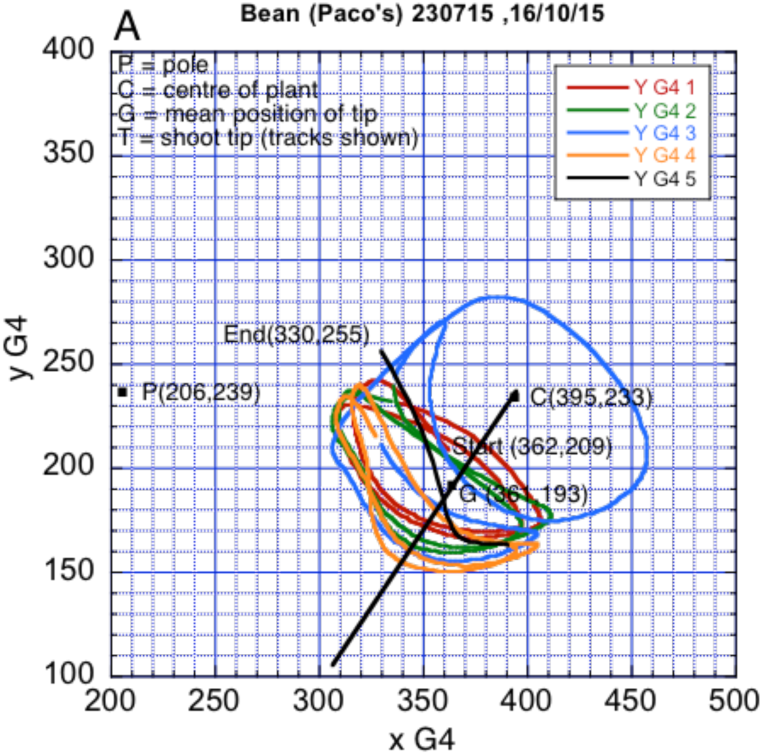
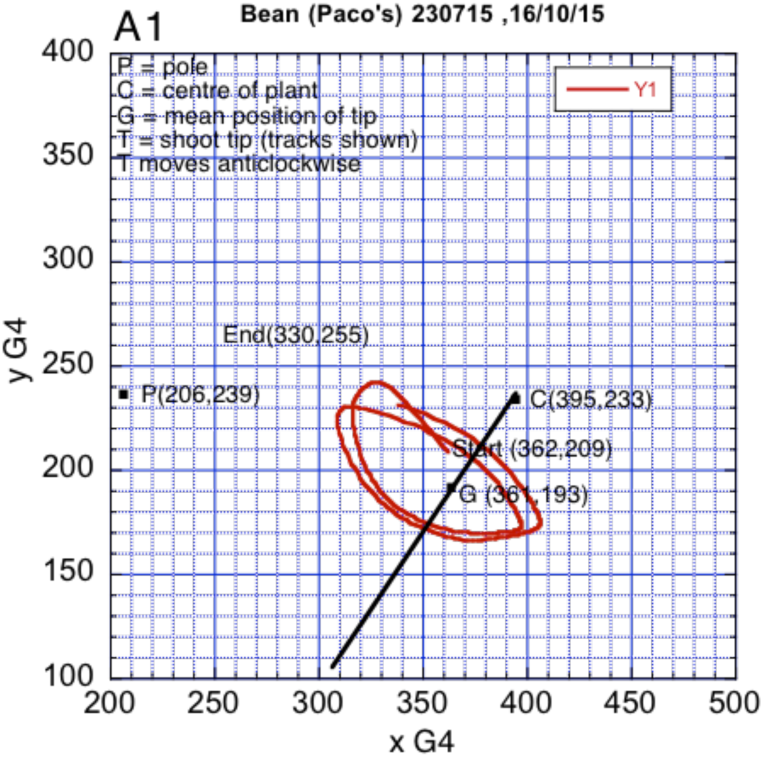
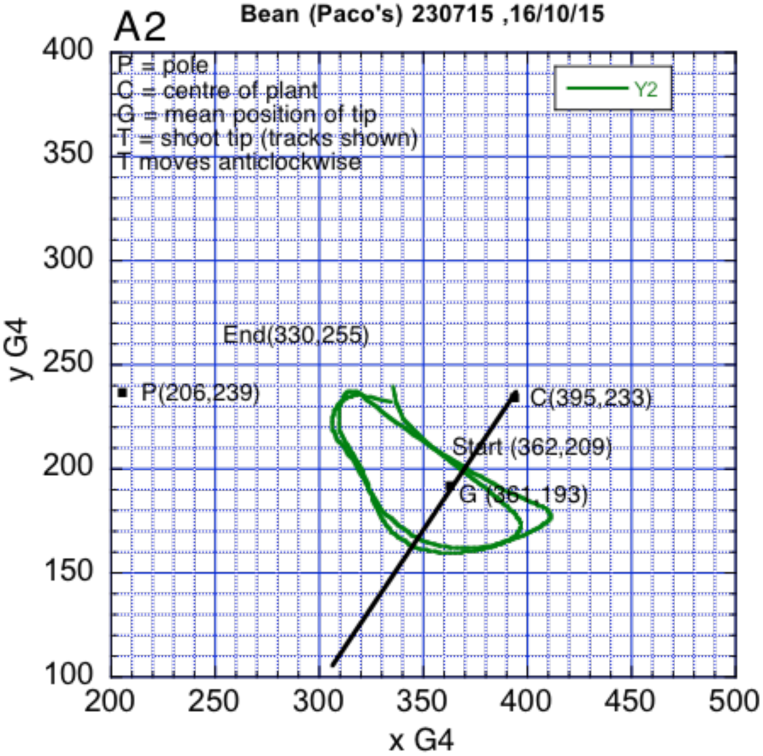
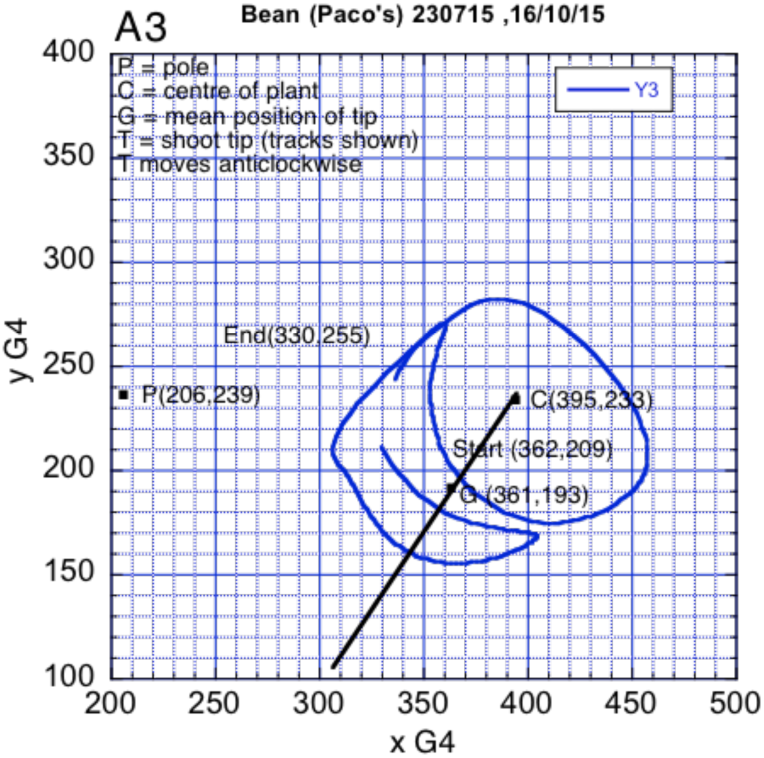
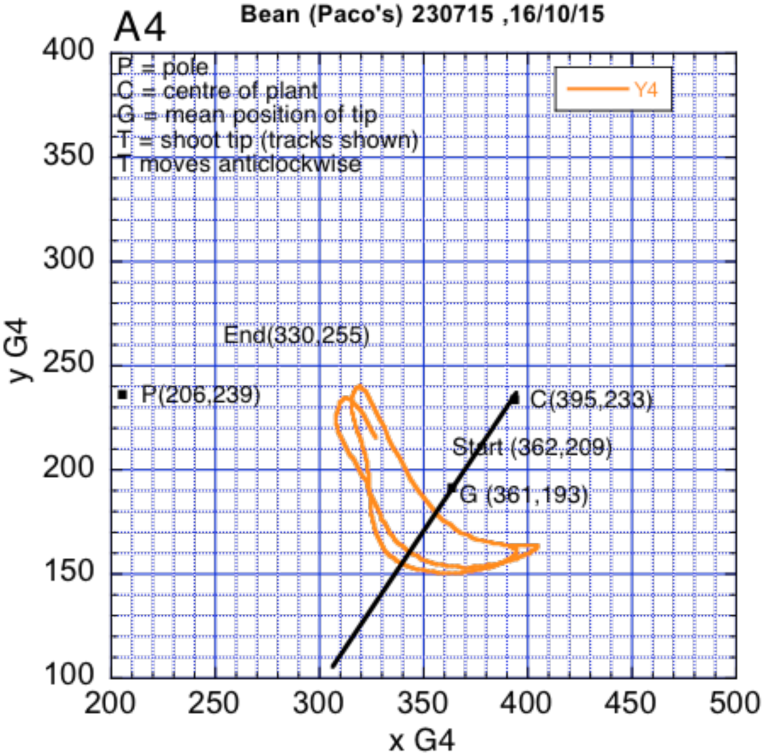
The (x,y) tracks of the tip of the shoot. The (x,y) coordinates were smoothed with a Gaussian filter sigma 4. This was the only smoothing done in the whole analysis. The colour coding (1 red, 2 green, 3 blue, 4 orange, 5 black) refers to four successive 0.5 video minute epochs from the start of the recording, ending with a 0.1 video minute epoch. (1 video minute = 25x60 = 1500 frames.

Figures A1-A4 below represent figure A split up into four successive 62.5 min epochs, to see whether there was any progressive shift in the orientation of the trajectory of T, or any progressive shift in the position of the trajectory along the CG line. There does not appear to be any such progressive shift in the orientation of the trajectory of T, but there appears to be some shift away from C (except for A3, which is an odd-ball).

Figure B plots, in green, against time, the radial distance from G to T (rGT) The mean strength (% of data fitting the theory with r^2^>0.95)) of tauG-guidance of rGT was high when rGT was decreasing, but significantly (p<0.01, t-test) lower when rGT was increasing (96.03% ± 0.56%se vs 85.43% ± 3.65%se). At the same time, the mean k value of tauG-guidance was significantly (p<0.01, t-test) *higher* when rGT was decreasing (0.55 ± 0.09se vs 0.29 ± 0.03se). These results indicate that the movement of the tip relative to its mean position (G) was very gently, but weakly tauG-guided when the tip was cast out (like a fly-fishing line) but was less gently, and strongly tauG-guided when it was being reeled in.

**Fig. B.**
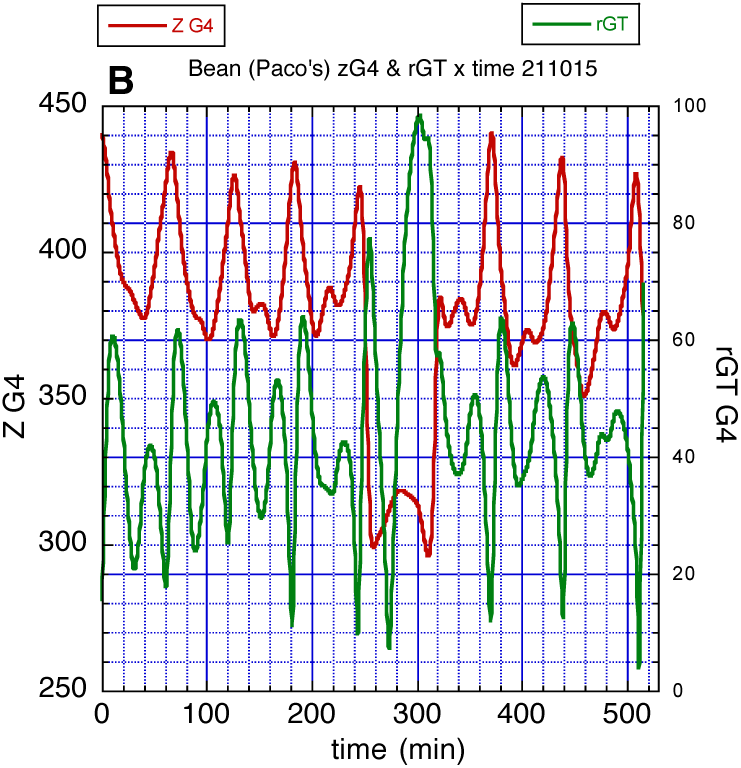
Plots against time of the radial distance from G to T, rGT (green) and the z coordinate of T (red).

Figure B also plots, in red, the vertical z coordinate of the shoot tip. Generally speaking, the tip tended to move down while it was being cast out, and move up while it was being reeled in (again like a fly-fishing line). Both up and down movements were strongly tauG-guided (up 95.48% ± 1.04%se; down 94.54% ± 0.49%se). The mean k values of 0.425 (up) and 0.356 (down) indicate gentle approach to both the zenith and bottom.

To obtain a measure of the angular speed of circumnutation, Figure C plots the angle, GTx, between GT and the x-axis. The angle increases continuously from 0 to 2670 degrees, at an approximately steady speed of 10.9 degrees per minute (r^2^ = 0.996), except for a sequence of small glitches.

**Figure C.**
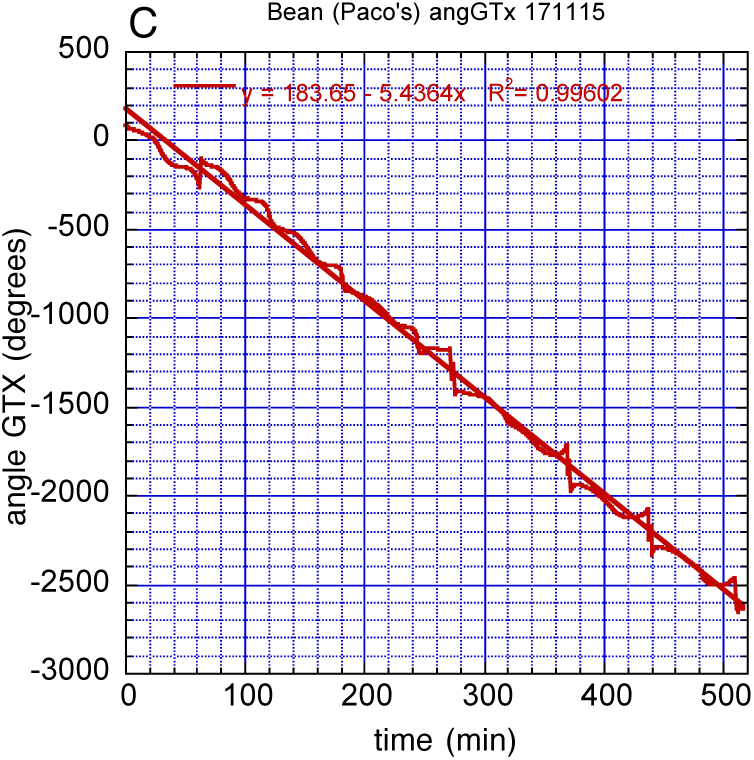
Plot of the angle GTx between GT and the x-axis.

Finally, Figure D plots the distance rTP between T and P. T was moderately strongly tauG-guided when receding from and approaching P (receding 92%; approaching 94%). However, it never reached the pole (see discussion section, below).

**Figure D.**
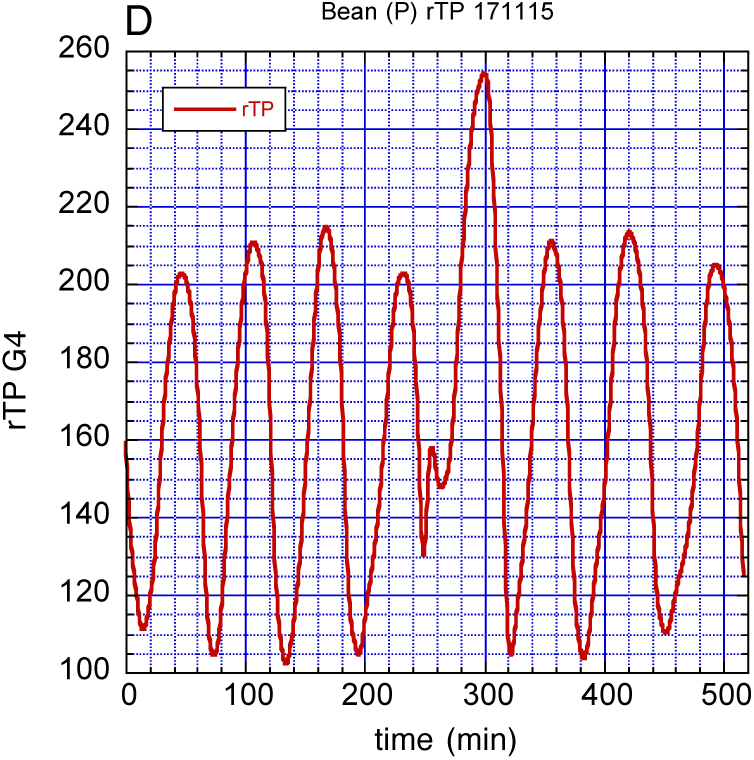
Plot of the distance rTP between T and P.

## Discussion

The results reported here are congruent with the working hypothesis that climbing plants resonate to specificational information of the type provided by high-level, relational invariants such as rho/tau—variables that guide interactions with the environment at an ecological scale. This provides grounds to argue plants perceive and circumnutate in their surroundings ecologically: they explore their environment by picking up invariant information in the form of relational properties that can be detected directly, unaided by any additional processing of information.

Nevertheless, even acknowledging the possibility that climbing plants may guide their movement of circumnutation ecologically, someone may resist the parallelism between plants and animals, wondering for example what sensory modality is involved in the direct process of perception, or what type of non-neural substrate could permit the detection of ecological information. After all, basic and cognitive neuroscience appears to have a pretty good story to tell both at the level of the sensory modalities involved in animal perception, as well as at the level of the neural correlates involved. But, how do neuron-less living organisms cope with their demands, such as perceiving an object as a potential support, despite lacking eyes or ears, and a nervous system at all?

To make a long story short, general rho/tau theory is neither (sensory) modality-specific nor substrate-specific (Calvo et al., 2014). Rho/tau related informational variables remain specificational regardless of the sensory modality involved. Consider for illustration a free-swimming cell (Delafield-Butt et al., 2013), where the tau of the cell is the time needed to swim to a cathode by sensing electric fields at its current rate-of-closing. Or take steering bats whose guidance is based on echolocation (Lee et al., 1995). It is thus not a unique sensory modality, such as vision, that drives the wheel, but rather the changes in the sensory gaps of *any sensory modality* whatsoever. They inform us as to which opportunities for behavioural output are available. As we saw earlier, one advantage of our ecological understanding of perception and action is that, unlike the size of motion-gaps, or any of its time derivatives (velocity, acceleration etc), rho of a motion-gap is, in principle, directly perceptible by all known perceptual systems. Given that plants have at their disposal a panoply of sensory modalities other than vision or hearing (Chamovitz, 2012) —in fact, plants can sense up to 22 different biotic and abiotic vectors (Trewavas, 2008), including electrical, magnetic, chemical, and vibrational fields—, plant perception and movement can be the subject to an ecological analysis in the very same way that a free-swimming cell and a bat are. Plant apexes contain electrical, chemical, vibrational, gravitational and optical sensory transducers that afford information about the movement of the apex in relation to the environment. Nevertheless, more work is needed, and the preliminary results of this report do not allow us to single out one particular sensory modality to be involved in the control of circumnutation.

On the other hand, general rho/tau theory is substrate-neutral. The fact that guidance is partly performed intrinsically, does not mean that it should be neurally based. Regardless of the type of substrate involved, what counts is whether the spatiotemporal scale of processes remains ecological or not. If specificational information happens to be found, for instance, at the scale of hormonal processes, the relevant substrate may then be hormonal.

With that being said, plants and animals share many ‘neural’ features (Calvo, submitted). A number of plant neurotransmitters have been identified (Baluška & Mancuso, 2009a). The role of G-aminobutyric acid (GABA) in plant signalling, for instance, is generating increasing interest (Bouché and Fromm, 2004). In fact, neuroid conduction (Mackie, 1970) is a basic and widespread form of signalling. It is well-known how electrical events propagate in the non-nervous cells of protists and plants. Plant and animal cells conduct signals from receptor to effector sites. Information is conveyed through an electro-chemical communication system (Keijzer et al., 2013), and action potentials propagate multidirectionally along the phloem (Baluška & Mancuso, 2009b), allowing plants to elaborate coordinated responses.

In multicellular animals, relevant information takes the form of temporal changes in ‘neural power’ (the rate of flow of electrochemical energy that flows along nerves, either continuously as a graded potential or as trains of action potentials). Prescriptive neural-power gaps are generated in the central nervous system. The analysis of skilled movement in animals has permitted identification of two types of prescriptive neural-power gaps: G-type and D-type, depending on whether the gap closes at constant acceleration or deceleration.

In cells, micro-organisms such as bacteria, and plants, this information-bearing ‘neural power’ consists in streams of ions flowing along ion channels, and thus renders itself subject to the same type of analysis. Unfortunately, because our recording started when the bean plant was already circumnutating, and ended when it died of heat exhaustion, before it had attempted to reach the support, we are unable to analysis whether the skilled movement of circumnutation performed by climbing plants conforms to prescriptive neural-power gaps of G-type or D-type in the control of the approach movement towards the support.

Climbing plants perform an ordinary movement of circumnutation in the early stages of development. As they grow the pattern of nutation changes, and here is where we suspect that such analysis would bear fruit. The sophistication of modified circumnutation is something that Darwin himself had already noticed. In a description of the circumnutation of *Ceropegia*, he observed:

> When a tall stick was placed so as to arrest the lower and rigid internodes of the *Ceropegia*, at the distance at first of 15 and then of 21 inches from the centre of revolution, the straight shoot slowly and gradually slid up the stick, so as to become more and more highly inclined, but did not pass over the summit. Then, after an interval sufficient to have allowed of a semi-revolution, the shoot suddenly bounded from the stick and fell over to the opposite side or point of the compass, and reassumed its previous slight inclination. It now recommenced revolving in its usual course, so that after a semi-revolution it again came into contact with the stick, again slid up it, and again bounded from it and fell over to the opposite side. This movement of the shoot had a very odd appearance, as if it were disgusted with its failure but was resolved to try again. (Darwin, 1875, pp. 12-13)

We have observed similar surprising ways in which climbing plants sway away from the regular elliptical revolution in a way that is congruent with the hypothesis that the bean perceives the support and tries again and again by elongating to reach it. Unfortunately, the videos where this movement is observed do not render themselves to a rho/tau analysis since the coordinates cannot be extracted and corrected with sufficient accuracy as to analyse them properly. One of our future objectives is to perform a rho/tau analysis of the whole pattern of movement, from original circumnutation all the way to the twining and securing of the support.

## Conclusion

In *The movements and habits of climbing plants* Darwin observes:

> It has often been vaguely asserted that plants are distinguished from animals by not having the power of movement. It should rather be said that plants acquire and display this power only when it is of some advantage to them; this being of comparatively rare occurrence, as they are affixed to the ground, and food is brought to them by the air and rain. (p. 206).

We are now aware not only that plants’ behaviour is reversible, non-automatic, and repeatable in a manner that responds to metabolically salient features of the environment (Calvo et al., 2014), but also of the increasing degree of sophistication of plant movements as a function of the specific goal to be attained. Anthony Trewavas considers the stilt palm (Allen, 1977), a plant whose light-foraging behaviour results in the selective growing of new roots in the direction of sunlight, letting the older ones die. Trewavas writes:

> the filiform stem explores, locates and recognizes a new trunk and reverses the growth pattern. As it climbs, the internode becomes progressively thicker and leaves progressively redevelop to full size…This behaviour is analogous to animals that climb trees to forage, intelligently descend when food is exhausted or competition severe, and then climb the next tree. (2003, p. 15)

Other strategies plants have evolved include the capacity to selectively becoming mobile or sessile, alternatively, as a function of the environmental demands (Ray, 1992). Twining around a support is certainly not the only pattern of movement of interest in the plant kingdom. We may thus wonder whether the ecological laws of animal goal-directed movement apply to plants more generally. In fact, further potential applications of general tau/rho theory to plants include analysing, for instance, how orchid flowers orient to gravity (with their ‘chins’ down); shoots grow up from an initial horizontal orientation; stems orient their flowers to light; root tips guide their growth downward; root tips guide their growth away from light; etc. Although more research is needed, the results reported here are consistent with an ecological interpretation of the power of movement of plants.

## Acknowledgments

This research was supported by Fundación Séneca-Agencia de Ciencia y Tecnología de la Región de Murcia, through project 11944/PHCS/09 to PC and DL. We thank Antonio Guirao for providing us with the perspective-error correction equations and figures 1 and 2.

## Author contributions

D.N.L. and P.C. conceived, designed the experiments, and wrote the paper. V.R. digitized the time-lapse frames. P.C. and V.R. prepared the experimental set up at MINT Lab, performed the experiments, and post-processed the data. D.N.L. performed the rho/tau analysis.

## Footnotes

1 The degree of rhoG-guidance of Y is assessed by linearly regressing the measured value, *ρ*(*Y,t*), on the mathematical function *ρ*(*G,t,T_G_*)(Schogler et al. 2008). The criterion used as evidence of rho-Gguidance is that more than 95% of the variance in the data is accounted for by Eq. (3) (i.e., r^2^ >0.95). When this criterion is not met for a whole movement, the maximum percentage of the data extending to the end of the movement that satisfies the criterion is computed. The regression slope measures *λ_Y,G_*.

2 The values of X, t, and TG have been normalized for clarity, without loss of generality in all figures: the normalized size of the gap X equals 1; the gap starts to close at normalized time –1 and ends closure at normalized time 0; the normalized duration of closure, TG, equals 1.

3 Optical distortion due to the webcam lens curvature was negligible and only parallax, perspective-induced error was corrected during post-processing

## References

Allen, P. H. (1977). The Rain Forests of Golfo Dulce. Stanford: Stanford University Press.

Baluška, F., & Mancuso, S. (2009a). Deep Evolutionary Origins of Neurobiology: Turning the essence of ‘neural’ upside-down. Communicative & Integrative Biology, 2(1): 60–65.

Baluška, F., & Mancuso, S. (2009b). Plants and Animals: Convergent evolution in action? In F. Baluška (Ed.), Plant-Environment Interactions: From Sensory Plant Biology to Active Plant Behavior (285–301). Berlin, Germany: Springer-Verlag.

Bouché, N., & Fromm, H. (2004). GABA in Plants: Just a metabolite? Trends in Plant Science, 9(3): 110–115.

Calvo, P. (2016). The Philosophy of Plant Neurobiology: A manifesto. Synthese 193: 1323–1343.

Calvo, P., & Baluška, F. (2015). Conditions for Minimal Intelligence across Eukaryota: A cognitive science perspective. Frontiers in Psychology, 6: 13–29. doi: 10.3389/fpsyg.2015.01329

Calvo, P., & Keijzer, F. (2011). Plants: Adaptive behavior, root brains, and minimal cognition. Adaptive Behavior, 19(3): 155–171.

Calvo, P., Martín, E., & Symons, J. (2014). The Emergence of Systematicity in Minimally Cognitive Agents. In P. Calvo, & J. Symons (Eds.), The Architecture of Cognition: Rethinking Fodor and Pylyshyn’s systematicity challenge (397–434). Cambridge, MA: MIT Press.

Carello, C., Vaz, D., Blau, J. J. C., & Petrusz, S. C. (2012). Unnerving intelligence. Ecological Psychology, 24(3): 241–264.

Chamovitz, D. (2012). What a Plant knows: A field guide to the senses. New York, NY: Scientific American/Farrar, Staus & Giroux.

Craig, C. M., Delay, D., Grealy, M. A., & Lee, D. N. (2000). Guiding the Swing in Golf Putting. Nature, 405: 295–296.

Craig, C. M., Lee, D. N. (1999). Neonatal Control of Nutritive Sucking Pressure: Evidence for an intrinsic tau-guide. Experimental Brain Research, 124: 371–382.

Darwin, C. (1875). The Movements and Habits of Climbing Plants. London, UK: John Murray.

Darwin, C., & Darwin, F. (1880). The Power of Movement in Plants. London, UK: John Murray.

Delafield-Butt, J. T., Galler, A., Schögler, B., & Lee, D. N. (2010). A Perception-action Strategy for Hummingbirds. Perception, 39: 1172–1174.

Delafield-Butt, J. T., Pepping, G-J., McCaig, C. D., & Lee, D. N. (2012). Prospective Guidance in a Free-swimming Cell. Biological Cybernetics, 106: 283–293.

Gibson, J. J. (1966). The Senses considered as Perceptual Systems. Boston: Houghton Mifflin.

Gibson, J. J. (1979). The Ecological Approach to Visual Perception. Boston: Houghton Mifflin.

Grealy, M. A., Craig, C. M., & Lee, D. N. (1999). Evidence for On-line Visual Guidance during Saccadic Gaze Shifts. Proceedings of the Royal Society of London B, 266: 1799–1804.

Isnard, S., & Silk, W. K. (2009). Moving with Climbing Plants from Charles Darwin’s Time into the 21st Century. American Journal of Botany, 96(7): 1205–1221.

Keijzer, F., van Duijn, M., & Lyon, P. (2013). What Nervous Systems do: Early evolution, input-output versus skin brain theory. Adaptive Behavior, 21(2): 67–85.

Lee, D. N. (1998). Guiding Movement by Coupling Taus. Ecological Psychology, 10: 221–250.

Lee, D. N. (2005). Tau in Action in Cevelopment. In J. J. Rieser, J. J. Lockman, & C. A., Nelson (Eds.), Action as an Organizer of Learning and Development (3–49). Hillsdale, NJ: Erlbaum.

Lee, D. N., Bootsma, R. J., Frost, B. J., Land, M., & Regan, D. (2009). General Tau Theory: Evolution to date. Special Issue: Landmarks in Perception. Perception, 38: 837–858.

Lee, D. N., Craig, C. M., Grealy, M. A. (1999). Sensory and Intrinsic Coordination of Movement. Proceedings of the Royal Society of London B, 266: 2029–2035.

Lee, D. N., Georgopoulos, A. P., Clark, M. J. O., Craig, C. M., & Port, N. L. (2001). Guiding Contact by Coupling the Taus of Gaps. Experimental Brain Research, 139: 151–159.

Loeb, J. (1918). Forced Movements, Tropisms, and Animal Conduct. Philadelphia and London: J.B. Lippincott.

Mackie, G. O. (1970). Neuroid Conduction and the Evolution of Conducting Tissues. The Quarterly Review of Biology, 45(4): 319–332.

Merchant, H., Battaglia-Mayer, A., & Georgopoulos, A. P. (2004). Neural Responses during Interception of Real and Apparent Circularly Moving Stimuli in Motor Cortex and Area 7a. Cerebral Cortex, 14: 314–331.

Mugnai, S., Azzarello, E., Masi, E., Pandolfi, C., & Mancusi, S. (2007). Nutation in Plants. In S. Mancuso, & S. Shabala (Eds.), Rhythms in Plants: Phenomenology, mechanisms, and adaptative significance (77–79). Berlin, Germany: Springer.

Padfield, G. D. (2011). The Tau of Flight Control. The Aeronautical Journal, 115(1171): 521–556.

Ray, T. S. (1992). Foraging Behaviour in Tropical Herbaceous Climbers (Araceae). Journal of Ecology, 80: 189–203.

Richardson, M., Shockley, K., Fajen, B. R., Riley, M. A., & Turvey, M. (2008). Ecological Psychology: Six Principles for an Embodied-Embedded Approach to Behavior. In P. Calvo, & T. Gomila, Handbook of Cognitive Science: An Embodied Approach (161-187). San Diego, USA: Elsevier.

Rind, F. C., & Simmons, P. J. (1999). Seeing what is coming: Building collision-sensitive neurons. Trends in Neurosciences, 22: 215–220.

Schogler, B., Pepping, G. J., Lee, D. N. (2008). TauG-guidance of Transients in Expressive Musical Performance. Experimental Brain Research, 198: 361–372.

Sun, H., Frost, J. F. (1998). Computation of different Optical Variables of Looming Objects in Pigeon nucleus rotundus Neurons. Nature Neuroscience, 1: 296–303.

Trewavas, A. J. (2003). Aspects of Plant Intelligence. Annals of Botany, 92: 1–20.

Trewavas, A. J. (2008). Aspects of Plant Intelligence: Convergence & Evolution. In S. C. Morris (Ed.), The Deep Structure of Biology: Is convergence sufficiently ubiquitous to give a directional signal? (68–110). West Conshohocken, PA: Templeton Press.

Turvey, M., Shaw, R., Reed, E. S., & Mace, W. (1981). Ecological Laws for Perceiving and Acting: A reply to Fodor and Pylyshyn. Cognition, 10: 237–304.

Van der Meer, A. L. H., Van der Weel, F. R., & Lee, D. N. (1994). Prospective Control in Catching by Infants. Perception, 23: 287–302.

Van der Weel, F. R., & Van der Meer, A. L. H. (2009). Seeing it coming: Infants’ brain responses to looming danger. Naturwissenschaften, 96: 1385–1391.

Wagner, H. (1982). Flow-field Variables trigger Landing in Flies. Nature, 297: 147–148.

